# Interaction of the mitochondrial calcium/proton exchanger TMBIM5 with MICU1

**DOI:** 10.1101/2025.03.28.645939

**Authors:** Li Zhang, Benjamin Gottschalk, Felicia Dietsche, Sara Bitar, Diones Bueno, Liliana Rojas-Charry, Anshu Kumari, Vivek Garg, Wolfgang F. Graier, Axel Methner

**Author notes:** Correspondence should be addressed to: Axel Methner MD, University Medical Center of the Johannes Gutenberg-University Mainz, Institute for Molecular Medicine, Langenbeckstr. 1, D-55131 Mainz, Germany.

## Abstract

Ion transport within mitochondria influences their structure, energy production, and cell death regulation. TMBIM5, a conserved calcium/proton exchanger in the inner mitochondrial membrane, contributes to mitochondrial structure, ATP synthesis, and apoptosis regulation. The relationship of TMBIM5 with the mitochondrial calcium uniporter complex formed by MCU, MICU1-3, and EMRE remains undefined. We generated *Tmbim5*-deficient Drosophila that exhibit disrupted cristae architecture, premature mitochondrial permeability transition pore opening, reduced calcium uptake, and mitochondrial swelling – resulting in impaired mobility and shortened lifespan. Crossing these with flies lacking mitochondrial calcium uniporter complex proteins was generally detrimental, but partial MICU1 depletion ameliorated the *Tmbim5*-deficiency phenotype. In human cells, MICU1 rescues morphological defects in TMBIM5-knockout mitochondria, while TMBIM5 overexpression exacerbates size reduction in MICU1-knockout mitochondria. Both proteins demonstrated opposing effects on submitochondrial localization and coexisted in the same macromolecular complex. Our findings establish a functional interplay between TMBIM5 and MICU1 in maintaining mitochondrial integrity, with implications for understanding calcium homeostasis mechanisms.

## Introduction

Mitochondria are intracellular organelles that harness a proton gradient generated by the electron transfer system in the inner mitochondrial membrane (IMM) to propel the F_0_F_1_-ATP synthase in a process called oxidative phosphorylation. The negative membrane potential also drives the passive uptake of Ca^2+^ ions, which adapts mitochondrial ATP generation to demand by matching cellular and mitochondrial Ca^2+^ levels during cellular activity and is important for cellular Ca^2+^ signaling (reviewed by (Giorgi *et al*, 2018)).

Mitochondrial Ca^2+^ uptake predominantly occurs through the mitochondrial Ca^2+^ uniporter (MCU), a pore-forming protein (Baughman *et al*, 2011; De Stefani *et al*, 2011). The MCU is the key component of the mitochondrial Ca^2+^ uniporter complex (MCUC), which includes several interacting and regulatory proteins such as MICU1 (Mitochondrial Calcium Uptake 1) (Mallilankaraman *et al*, 2012), MICU2 (Patron *et al*, 2014), MICU3 (Patron *et al*, 2019), the Essential MCU Regulator (EMRE) (Vais *et al*, 2016; Sancak *et al*, 2013), and a dominant-negative subunit called MCUb (Raffaello *et al*, 2013). Despite its critical role, the knockout (KO) of *Mcu* in mice exhibits strain-dependent effects, ranging from a mild skeletal muscle phenotype (Pan *et al*, 2013) to lethality during late embryonic development (Luongo *et al*, 2015). These observations suggest the presence of alternative mechanisms for Ca^2+^ entry into mitochondria (Murphy *et al*, 2014; Wüst *et al*, 2017).

Recently, others and we proposed that the ubiquitously expressed protein Transmembrane Bax-Inhibitor Motif (TMBIM)-containing protein 5 (TMBIM5) represents an additional Ca^2+^ and proton exchange system in the IMM. Austin et al. (Austin *et al*, 2022) and Patron et al. (Patron *et al*, 2022) both proposed that TMBIM5 works as an outward Ca^2+^ transport system while we observed increased mitochondrial Ca^2+^ uptake in cells overexpressing TMBIM5 stimulated with ER Ca^2+^ release agents but no changes in mitochondrial release (Zhang *et al*, 2022). These results suggest that this new transport system can work in both directions depending on cellular demands. TMBIM5 is also part of the MCUC interactome as shown by proteomics (de la Herran *et al*, 2024), but the functional crosstalk between TMBIM5 and the MCUC is not well defined. This represents a significant gap in our understanding of the integrated networks regulating mitochondrial Ca^2+^ homeostasis, cristae architecture, and tissue-specific vulnerabilities to mitochondrial dysfunction.

Mutation of the proposed channel pore of TMBIM5 in mice caused a skeletal myopathy associated with swollen mitochondria and a disrupted cristae architecture (Zhang *et al*, 2022). Also in cells, *TMBIM5* deficiency leads to a disrupted cristae architecture (Seitaj *et al*, 2020; Oka *et al*, 2008) and an increased release of cytochrome *c* in response to apoptotic stimuli (Oka *et al*, 2008). Additionally, TMBIM5 associates with CHCHD2 (Coiled-Coil-Helix-Coiled-Coil-Helix Domain Containing 2), a protein that is part of the Mitochondrial Contact Site and Cristae Organizing System (MICOS) complex (Meng *et al*, 2017). CHCHD2 deficiency results in similar effects as TMBIM5 deficiency, including altered cristae architecture and increased cytochrome *c* release (Meng *et al*, 2017). This is of interest because mutations in the *CHCHD2* gene are linked to autosomal-dominant familial Parkinson’s disease (Funayama *et al*, 2015). Together these findings suggest that TMBIM5 is involved not only in Ca^2+^ transport but also in the stabilization of cristae architecture and the regulation of cytochrome *c* release.

Similar to *Tmbim5* loss of function in mice, mutations of *MICU1* in humans cause a proximal myopathy additionally associated with learning difficulties and a progressive extrapyramidal movement disorder (Logan *et al*, 2014). A detailed analysis of the mitochondrial Ca^2+^ uptake phenotype associated with disease-causing mutations and a skeletal muscle-specific knockout of *Micu1* in mice demonstrated a lower threshold for MCU-mediated Ca^2+^ uptake but also impaired mitochondrial Ca^2+^ uptake during excitation-contraction, aerobic metabolism impairment, muscle weakness, fatigue, and myofiber damage during physical activity (Debattisti *et al*, 2019). Also similar to TMBIM5, MICU1 appears to play a role in the stability of cristae junctions and cytochrome *c* release (Gottschalk *et al*, 2019) and interact with CHCHD2 (Tomar *et al*, 2023). Despite these parallel phenotypic consequences and shared interaction partners, a potential functional relationship between TMBIM5 and MICU1, remains inadequately characterized.

Besides a direct interaction also a more indirect interplay between TMBIM5 and the MCUC appears possible because TMBIM5 interacts with and inhibits AFG3L2 (Patron *et al*, 2022). AFG3L2 forms part of the m-AAA protease complex, which degrades misfolded or damaged proteins within the IMM. Importantly, the m-AAA protease also degrades non-assembled EMRE, the above-mentioned essential MCU regulator. This serves to ensure the efficient assembly of gatekeeper subunits with MCU and prevents the accumulation of constitutively active MCU-EMRE channels (König *et al*, 2016; Tsai *et al*, 2017). Changes in AFG3L2 activity imposed by changes in TMBIM5 abundance or interaction could, therefore, indirectly affect the composition of the MCUC. EMRE was not identified in the proteome analysis of TMBIM5-deficient cells reported by Patron et al. (Patron *et al*, 2022).

A recent comprehensive analysis of the phenotype of MCUC deficiency in Drosophila melanogaster strengthened the assumption that MICU1 has roles beyond regulating Ca^2+^ entry through the MCUC (Tufi *et al*, 2019). The fly genome encodes MCU, EMRE, MICU1, and MICU3, but lacks MICU2 and MCUb. Knockout of Mcu or Emre abolishes rapid mitochondrial Ca^2+^ uptake but results in only mild phenotypes and slightly shortened lifespans. In contrast, loss of Micu1 is developmentally lethal and this cannot be rescued by simultaneous knockout of Mcu or Emre, indicating again uniporter-independent functions for Micu1 as also mentioned above. Mutants for Micu3 are viable but exhibit mild neurological impairments, and Micu1 and Micu3 are not functionally interchangeable (Tufi *et al*, 2019).

In our study, we demonstrate that *Tmbim5* knockout in Drosophila closely resembles the findings obtained in mice. Genetic interaction studies with the MCUC proteins conserved in flies revealed that concomitant *Micu1* heterozygosity and knockdown mitigated the *Tmbim5* loss-of-function phenotype while concurrent knockout of the other MCUC components present in flies aggravated the phenotype. Further studies in human cells revealed a reciprocal regulatory relationship between MICU1 and TMBIM5 in mitochondrial structure maintenance. Our findings point to a critical functional interdependence between these proteins that maintains mitochondrial structural integrity, providing new insights into the molecular mechanisms governing Ca^2+^ homeostasis within mitochondria.

## Results

### TMBIM5 is upregulated in *MCU* and *MICU1* knockout cells

TMBIM5 is a Ca^2+^/H^+^ exchanger in the inner mitochondrial membrane (Patron *et al*, 2022; Austin *et al*, 2022; Zhang *et al*, 2022) and interacts with the MCUC (de la Herran *et al*, 2024). To investigate any potential compensatory changes of MCUC proteins in the absence of TMBIM5 hinting towards an involvement in common processes, we investigated the expression of MCU, MICU1, MICU2, and EMRE in *TMBIM5* knockout (KO) human embryonic kidney cells using immunoblotting. *TMBIM5* KO cells have a frameshift mutation (Figure 1A) introduced by CRISPR/Cas9-mediated gene editing (Zhang *et al*, 2022), resulting in loss of protein expression (Figure 1B). We found a slight reduction in MCU abundance in these *TMBIM5* KO cells (Figure 1C) but no changes in MICU1, MICU2 (Figure 1C) or EMRE (Figure 1D). The opposite experiment, studying TMBIM5 levels in the absence of MCUC proteins, demonstrated no apparent changes in *EMRE* KO cells (Figure 1D) but an increased abundance of TMBIM5 in *MCU* and *MICU1* but not *MICU2* KO cells (Figure 1E). We interpreted these results as indicative of a potential crosstalk between TMBIM5 and the MCUC, specifically MCU and MICU1. Importantly, EMRE levels were unchanged in *TMBIM5* KO and vice versa despite the inhibitory effect of TMBIM5 on the mAAA protease AFG3L2 (Patron *et al*, 2022), which controls EMRE levels (König *et al*, 2016).

**Figure 1.**
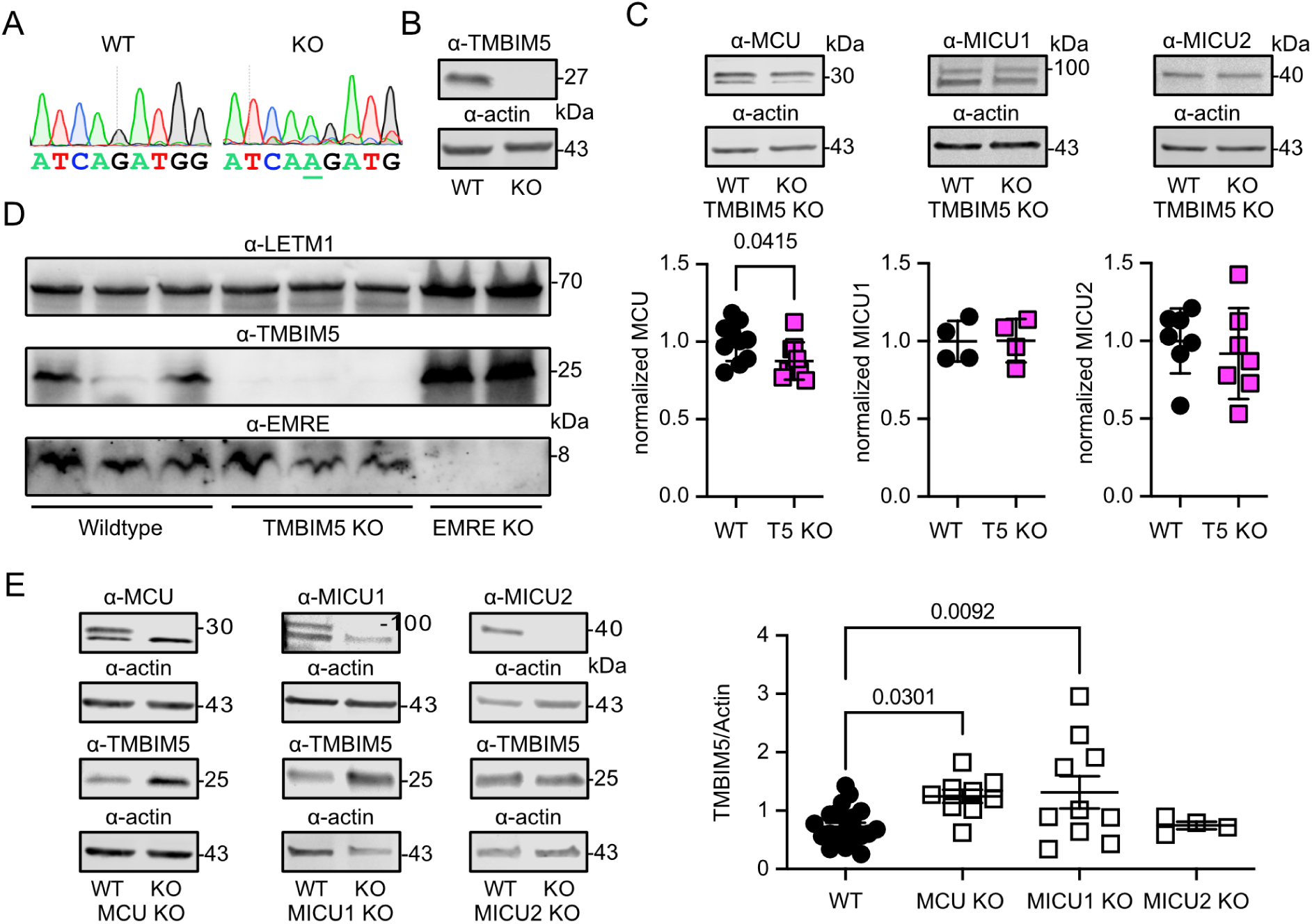
TMBIM5 is upregulated in MCU and MICU1 knockout cells. **(A)** DNA sequencing chromatograms of the genetic locus in wild type (WT) and TMBIM5-knockout (KO) HEK293 cells. The underlined additional adenosine results in a frameshift. **(B)** Immunoblot demonstrating loss of TMBIM5 protein expression in KO cells. Actin served as loading control, size is indicated. **(C)** Immunoblots demonstrating decreased levels of MCU (upper band, see E) and unchanged levels of MICU1 and MICU2 in TMBIM5 (T5) KO cells. Actin served as loading control, size is indicated. **(D)** Immunoblot demonstrating preserved EMRE protein levels in TMBIM5 KO cells. LETM1 served as loading control and EMRE KO cells to prove the specificity of the EMRE antibody. Size is indicated. **(E)** Immunoblots to demonstrate increased abundance of TMBIM5 in *MCU* and *MICU1* but not *MICU2* KO HEK293 cells. Actin served as loading control, size is indicated. Data in C and E are shown as mean±SD, each data point represents an independent immunoblot. Statistical significance was calculated by the student’s *t* test in C and one-way ANOVA followed by Holm-Šídák’s multiple comparisons test in E, *p* values are indicated.

### Drosophila *Tmbim5* knockout impairs wing posture, mobility, and lifespan

To investigate genetic interactions between TMBIM5 and MCUC proteins, we next generated and characterized a *Tmbim5*-deficient Drosophila melanogaster model. This approach was chosen for its efficiency and the availability of well-characterized fly lines with deletions in key MCUC components (Tufi *et al*, 2019). The fly genome encodes orthologs of MCU, EMRE, MICU1, and MICU3, but lacks MICU2 and MCUb. However, unlike vertebrates, flies have two mitochondrial TMBIM family members, *CG2076* and *CG1287*/*Mics1*. We previously demonstrated that ubiquitous *CG2076* knockdown causes pupal lethality, while *CG1287*/*Mics1* knockdown leads only to male sterility, consistent with its testis-specific expression (Zhang *et al*, 2021). These findings suggested that *CG2076* is the likely ortholog of TMBIM5 in Drosophila. We, therefore, now generated a complete knockout of *Tmbim5* in *w^1118^*flies using CRISPR/Cas9 technology (Figure S1) to study the effect of *Tmbim5* deficiency.

Because *Tmbim5* is located on the sex chromosome, all male flies have a complete knockout of *Tmbim5* (Figure 2A). Knockout males exhibit a wing posture phenotype suggestive of flight muscle degeneration (Figure 2B). This phenotype is often observed in flies lacking functional mitochondria (Clark *et al*, 2006; Park *et al*, 2006). In line with the knockdown results, *Tmbim5*-deficient flies also exhibit a severe reduction in their climbing ability (Figure 2C) and lifespan (Figure 2D), implying a strong effect of *Tmbim5* deficiency on the well-being of adult flies.

**Figure 2.**
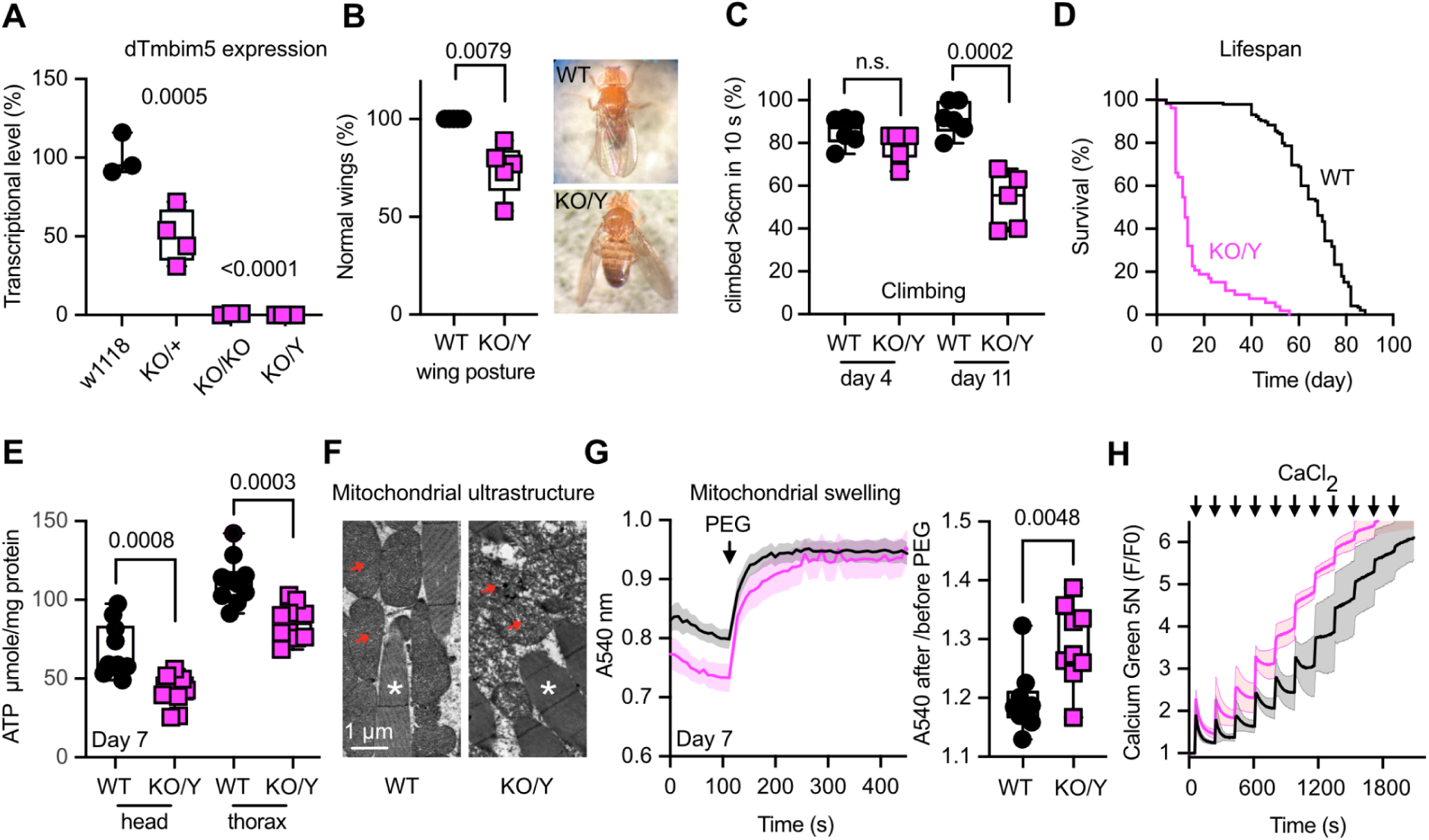
Drosophila *Tmbim5* knockout impairs wing posture, mobility, and lifespan with phenotypes resembling mammalian models: altered cristae, mitochondrial swelling, and reduced Ca²⁺ buffering capacity. **(A)** Tmbim5 expression assessed by quantitative PCR in wild type (WT, w^1118^), heterozygous (KO/+), homozygous (KO/KO), and hemizygous (KO/Y) knockout flies demonstrating loss of expression. Each data point contains five flies. **(B)** Abnormal wing posture in 6-week-old male KO/Y flies. Each data point contains the mean percentage of flies with normal wings in a sample of 25 flies. **(C)** Climbing ability of WT and KO/Y flies on the indicated days post-eclosion with an age-dependent decline in KO/Y. Each data point contains the mean speed of 25 flies. **(D)** Reduced lifespan of KO/Y flies, sample size >100 flies/group. **(E)** Ubiquitously reduced ATP levels in male *Tmbim5* knockout flies (KO/Y). Values were normalized to protein content. **(F)** Exemplary transmission electron microscopy images showing a disrupted cristae architecture of flight muscle mitochondria in KO/Y flies. Red arrows indicate mitochondria, asterisks mark indirect flight muscle. Size is indicated. **(G)** Larger delta in absorbance following osmotic shrinking with polyethylene glycol (PEG, added where indicated) in Tmbim5 KO mitochondria indicating swelling. Values were normalized to the last values after PEG-addition. Baseline absorbance and the ratio of absorbance before and after shrinking are quantified on the right. Data are shown as mean±SEM, *n*=9. **(H)** Reduced mitochondrial calcium buffering capacity in KO/Y flies shown by quantifying Ca^2+^ uptake of isolated mitochondria in a bath with Calcium Green 5N challenged by sequential addition of 5 µM CaCl_2_ pulses. Fluorescence was normalized to the initial value (F_0_). Data are shown as mean±SEM, *n*=5-9. Data are represented as box and whisker plots with the box representing the interquartile range, spanning from the 25th to the 75th percentile. A horizontal line within the box indicates the median. Whiskers extend from the minimum to the maximum data points. Each data point in A represented the mean value of 5 flies, in B, C 25 flies, in E 2 flies, in G and H 100 flies. Statistical significance was calculated by Šídák’s multiple comparisons test in A, the Mann Whitney test in B, unpaired *t* tests in C, E, G. *p* values are indicated.

### Mitochondrial phenotype of *Tmbim5* knockout resembles mammalian models: altered cristae, mitochondrial swelling, and reduced Ca²**⁺** buffering capacity

We then characterized the phenotype of these flies with a specific focus on mitochondrial function. Already at day 7 after eclosion, *Tmbim5* KO males have lower ATP levels in all major body parts (Figure 2E). This is not caused by a reduction in total mitochondrial mass quantified by the mRNA ratio of the mitochondrial and nuclear-encoded proteins of complex IV (Figure S2A) and quantification of citrate synthase, an enzyme in the tricarboxylic acid cycle (Figure S2B).

The cristae architecture in flight muscles was severely disrupted (Figure 2F). In embryonic kidney cells and mouse skeletal muscle this phenotype correlated with the observation that mitochondria were already swollen at steady state. We tested whether this is also the case in Drosophila *Tmbim5* KO mitochondria by adding polyethylene glycol to a suspension of mitochondria prepared from whole adult flies. This treatment osmotically shrinks the mitochondria to their minimum size (Luongo *et al*, 2017). To compare the starting baseline, we normalized the absorbance to the value after PEG addition. This revealed that *Tmbim5* KO mitochondria are indeed pre-swollen (Figure 2G), similar to mammalian TMBIM5 KO models. We next tested the total mitochondrial Ca^2+^ retention capacity by adding repetitive pulses of Ca^2+^ to isolated mitochondria. When the mitochondrial Ca²⁺ uptake capacity exceeds the organelle’s buffering capacity, excessive Ca²⁺ accumulation can lead to the opening of the permeability transition pore (PTP), a non-selective high-conductance pore that dissipates mitochondrial membrane potential. This demonstrated a faster opening of the mPTP in *Tmbim5* KO mitochondria (Figure 2H), suggesting an enhanced susceptibility of mutant mitochondria to this insult in line with the increased susceptibility to apoptosis and cytochrome *c* release in TMBIM5 loss-of-function cell models (Oka *et al*, 2008; Seitaj *et al*, 2020). Together, these results imply that *Tmbim5* deficiency in flies results in the same pathophysiological changes observed in mammalian systems. *Tmbim5*-deficient flies can thus serve as a *bona fide* model of TMBIM5 loss of function.

### Concomitant reduction of Micu1 mitigates lifespan reduction and attenuates mitochondrial defects in Tmbim5 knockout flies

To investigate potential compensatory changes of the MCUC similar to what we observed in *TMBIM5*-deficient HEK293 cells, we then studied the expression levels of *Tmbim5* in flies deficient in *Mcu*, *Emre*, *Micu1*, and *Micu3* (Tufi *et al*, 2019) and vice versa. This revealed an upregulation of *Mcu* mRNA in male *Tmbim5* KO flies but no changes in flies deficient in mRNA encoding the interacting proteins (Figure S3A). This is in contrast to the slight downregulation of MCU protein in human *TMBIM5* KO cells (Figure 1C), suggesting species-dependent differences or a post-translational regulation. We could not evaluate Tmbim5 protein levels in flies because it is not recognized by our antibodies. Interestingly, *Tmbim5* mRNA levels were also upregulated in *Mcu*-deficient flies, similar to *Mcu* being upregulated in *Tmbim5*-deficient flies; again, all other MCUC-deficient flies exhibited no statistically significant changes (Figure S3B and C).

To define generic interactions between the MCUC and Tmbim5, we next crossed *Tmbim5* KO males with MCUC-deficient flies. This drastically shortened the lifespan of double-deficient flies for *Mcu*, *Emre*, and *Micu3*-deficient flies (Figure 3A). However, the reduction of *Micu1* levels in *Micu1+/-* flies significantly increased the lifespan of wildtype and *Tmbim5* KO flies (Figure 3A) and completely normalized the detrimental effect of *Tmbim5* deficiency on climbing speed (Figure 3B). This was also true when we used RNAi-mediated knockdown to reduce *Micu1* in *Tmbim5* KO flies (Figure 3B). In line with the beneficial effect of *Micu1* depletion on lifespan and climbing speed of *Tmbim5*-deficient flies, *Micu1* heterozygosity also attenuated the effects of *Tmbim5* KO on faster mPTP opening (Figure 3C) and mitochondrial swelling (Figure 3D) and rescued the mitochondrial defects detected by transmission electron microscopy (Figure 3E). Together, these results imply that Micu1 reduction is largely beneficial and capable of mitigating the detrimental effects of Tmbim5 deficiency.

**Figure 3.**
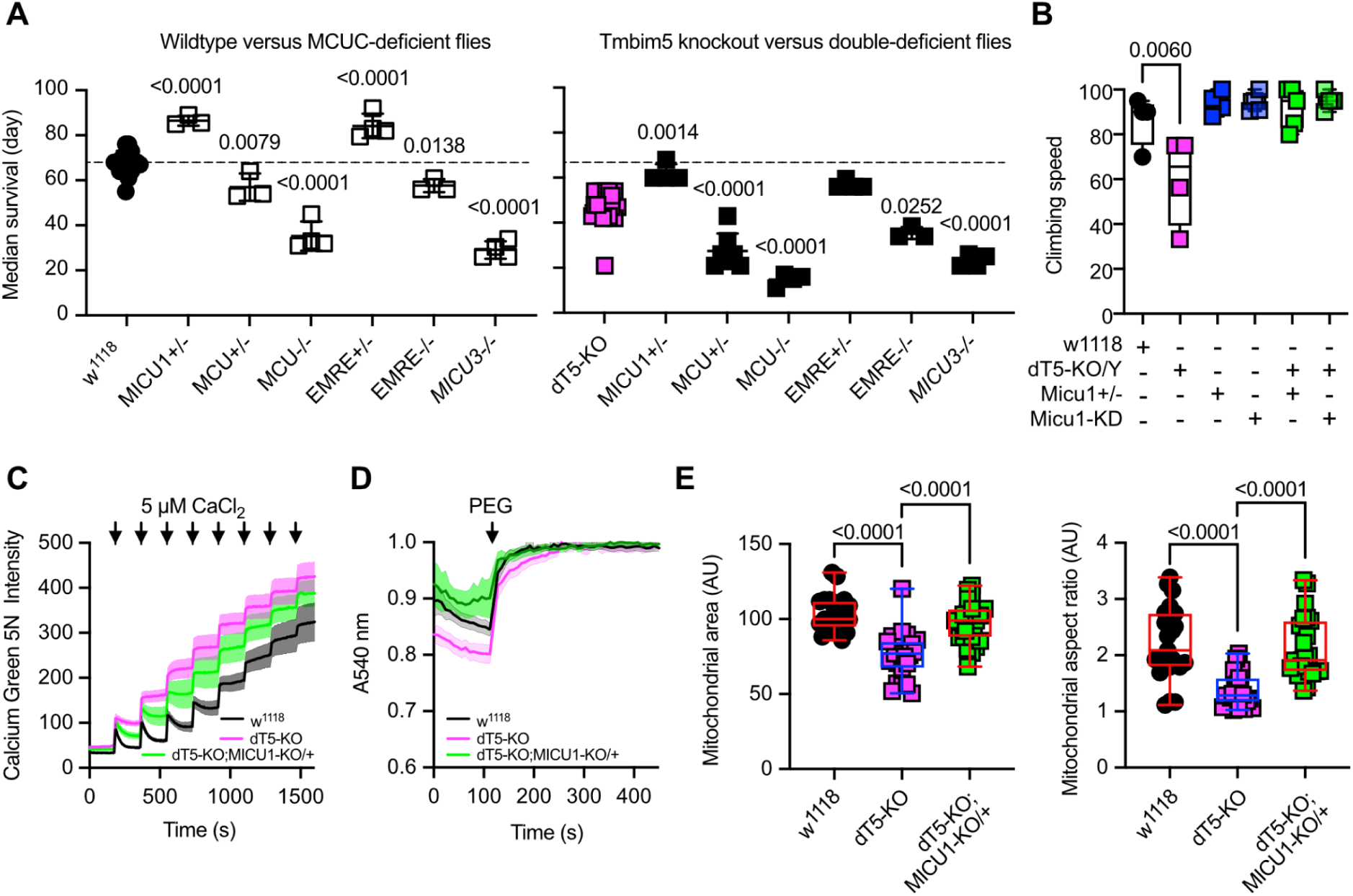
Concomitant reduction of Micu1 mitigates lifespan reduction and attenuates mitochondrial defects in Tmbim5 knockout flies. (A) Median survival of wild type (*w^1118^*), *Tmbim5* knockout (*dT5-KO/Y*), MCUC-deficient flies compared with flies with a combined deficiency as indicated. DKO, double knockout. Double deficiency was detrimental for *Mcu*, *Emre*, *Micu3* and beneficial for *Micu1*. (B) Climbing speed of *w^1118^*(wild-type), *dTmbim5* knockout (*dT5-KO/Y*), with (+) or without (−) heterozygous *Micu1* knockout (*Micu1+/-*), or ubiquitous *Micu1* knockdown using the GAL4-UAS system (*da-GAL4>UAS-Micu1-RNAi*, abbreviated as *Micu1-KD*). (C) Improved mitochondrial calcium buffering capacity in *dT5 KO*/*Micu1*+/-double-deficient flies shown by quantifying Ca^2+^ uptake of isolated mitochondria in a bath with Calcium Green 5N challenged by sequential addition of 5 µM CaCl_2_ pulses. Fluorescence was normalized to the initial value (F_0_). Data are shown as mean±SEM, *n*=5-9. (D) Reduced swelling of d*T5* KO/*Micu1*+/-double-deficient mitochondria shown by changes in absorbance following osmotic shrinking with polyethylene glycol (PEG, added where indicated). Values were normalized to the last values after PEG-addition. Baseline absorbance and the ratio of absorbance before and after shrinking are quantified on the right. Data are shown as mean±SEM, *n*=9. Each n represents a mitochondrial sample extracted from 100 flies. (E) Mitochondrial area and aspect ratio of *w^1118^*, *dT5-KO*, and *dT5-KO;Micu1+/-* flies detected by transmission electron microscopy and analyzed by ImageJ. Typical pictures are shown in Figure 2F. Data in panel A, B, E are presented as box and whisker plots with the box representing the interquartile range, spanning from the 25th to the 75th percentile. A horizontal line within the box indicates the median. Whiskers extend from the minimum to the maximum data points. Each data point in A and B represents the mean value of 25 flies. Statistical significance was determined using one-way ANOVA with Holm-Šídák’s multiple comparisons test in A, B and E, *p* values are indicated.

### TMBIM5 and MICU1 are found in the same macromolecular complex

To clarify whether the observed genetic interaction between MICU1 and TMBIM5 is mediated via physical proximity, we next analyzed the presence of TMBIM5 in macromolecular complexes containing MCU and MICU1 and assessed whether these complexes exhibit altered characteristics in the absence of TMBIM5. For this analysis, we utilized human cell models due to the availability of specific antibodies. We examined two human cell lines in which we previously generated and characterized *TMBIM5* knockout: HAP1 (Seitaj *et al*, 2020) and HEK293 (Zhang *et al*, 2022). Blue native gel electrophoresis revealed that in lysates from both cell lines, TMBIM5 and MICU1 predominantly migrated in a complex of approximately 140 kDa, with minimal presence in a larger 480 kDa complex containing MCU. Notably, TMBIM5 deficiency did not alter the size or stability of MCU- or MICU1-containing complexes (Figure 4A). These findings suggest the co-existence of MICU1 and TMBIM5 within a shared macromolecular complex. Consistent with this observation, we successfully co-immunoprecipitated native TMBIM5 with HA-tagged MICU1 but not with empty vector controls (Figure 4B). Reciprocal co-immunoprecipitation experiments using native MICU1 and TMBIM5 were unsuccessful, likely due to antibody limitations. Collectively, these results indicate that MICU1 and TMBIM5 may exist in the same macromolecular complex and interact under certain conditions, providing a potential molecular basis for their observed genetic relationship.

**Figure 4.**
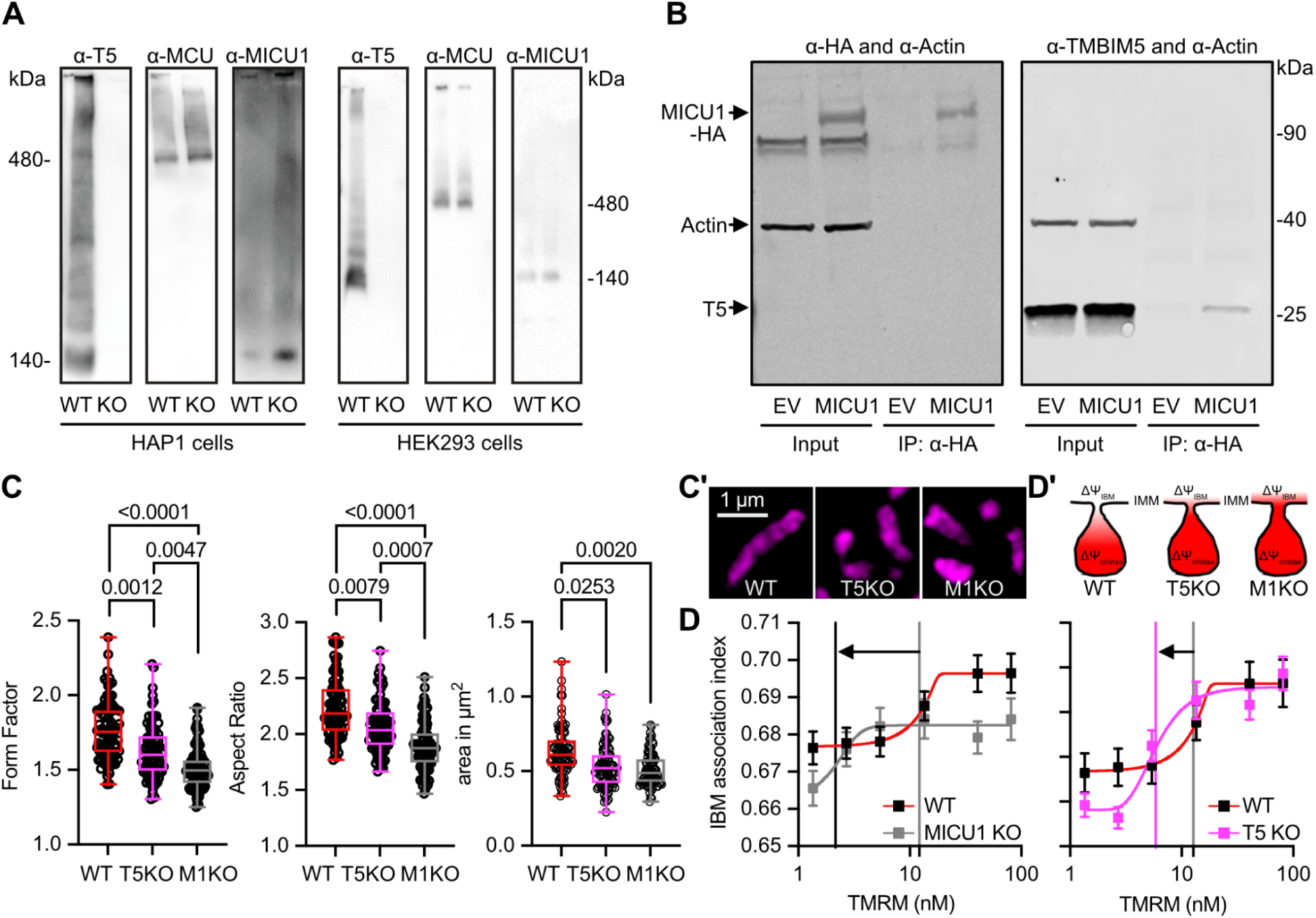
TMBIM5 and MICU1 are found in the same macromolecular complex and their deficiency results in similar changes in mitochondrial shape and inner boundary membrane. **(A)** Blue native gel electrophoresis of TMBIM5, MCU and MICU1 in human HAP1 (left) and HEK293 cells with (wild type, WT) and without TMBIM5 (knockout, KO) expression. Approximate molecular weight is indicated. **(B)** Co-immunoprecipitation of TMBIM5 with HA-tagged MICU1 (arrow). Left panel shows successful pull-down of tagged MICU1 and the right panel coimmunoprecipitation of endogenous TMBIM5. Molecular weight is indicated. **(C/C’)** Morphometric analysis of TMRM-labeled mitochondria in WT, TMBIM5 (T5) KO, and MICU1 (M1) KO HEK cells imaged with structured illumination microscopy. C’ shows typical pictures, size is indicated. **(D/D’)** Evaluation of mitochondrial membrane potential (ΔΨm) gradients of WT, T5KO, and M1KO HEK cells using structured illumination microscopy. D’ illustrates the findings. Data in C are presented as box and whisker plots with the box representing the interquartile range, spanning from the 25th to the 75th percentile. A horizontal line within the box indicates the median. Whiskers extend from the minimum to the maximum data points. Each data point represents a single cell accumulated from 4 independent experiments. Data in D show the mean ± standard errors of mean. Statistical significance in C was determined using the Kruskal-Wallis test followed by Dunn’s multiple comparisons test, *p* values are indicated.

### TMBIM5 and MICU1 deficiency results in similar changes in mitochondrial shape, size, and membrane potential compartmentalization

We next studied the effects of TMBIM5 and MICU1 deficiency on mitochondrial shape and size and found that both result in more rounded and smaller mitochondria with MICU1 knockout cells being even less tubular than TMBIM5 knockout cells while maintaining a similar size (Figure 4C). Besides its role as being the gatekeeper for the MCUC, preventing Ca^2+^ uptake at low cytosolic Ca^2+^ concentrations while facilitating uptake when Ca^2+^ levels rise above threshold (Mallilankaraman *et al*, 2012), MICU1 also independently functions as a structural component of the inner mitochondrial membrane (Gottschalk *et al*, 2019; Tomar *et al*, 2023). This membrane is structurally and functionally compartmentalized into two distinct domains: the cristae membrane (CM) and the inner boundary membrane (IBM). The IBM runs parallel to the outer mitochondrial membrane, maintaining a consistent distance and creating the intermembrane space, while the CM forms invaginations that project into the mitochondrial matrix. These domains are connected by narrow tubular structures called cristae junctions. MICU1 independently functions as a structural component of the IBM, where its exclusive localization through electrostatic interactions with cardiolipin stabilizes cristae junctions, maintains mitochondrial membrane potential, and prevents cytochrome c redistribution from cristae to the intermembrane space (Gottschalk *et al*, 2019). TMBIM5 is essential for maintaining cristae architecture (Oka *et al*, 2008; Seitaj *et al*, 2020; Zhang *et al*, 2022) and preventing cytochrome c release (Oka *et al*, 2008). Both proteins also interact with CHCHD2 (Meng *et al*, 2017; Tomar *et al*, 2023), a component of the MICOS. We, therefore, figured that they most probably functionally interact in their role in the maintenance of the IBM. We, therefore, quantified the IBM association index plotted against descending concentrations of TMRM in wildtype, TMBIM5 KO, and MICU1 KO cells to determine the relative membrane potential distribution between IBM and CM. This was achieved by using structured illumination microscopy, which allows the visualization and precise localization of TMRM within sub-mitochondrial compartments, and thus a visualization between the highly stained, highly polarized CM and lower stained, lower polarized IBM (Gottschalk *et al*, 2024). A higher IBM association index indicates relatively more TMRM accumulation in the IBM compared to the highly stained CM, reflecting a homogenization in membrane potential between these two compartments. Using this analysis, we again observed similar changes in TMBIM5 and MICU1 KO cells that were again more pronounced in MICU1-deficient cells (Figure 4D). A deficiency of both proteins resulted in a left shift of the IBM association index, indicating compromised electrochemical compartmentalization between mitochondrial subdomains (Figure 4D). This phenomenon reflects impaired maintenance of the membrane potential gradient that normally exists between the IBM and CM. Together, this implies that TMBIM5 and MICU1 functionally converge in maintaining mitochondrial shape, size, and membrane potential compartmentalization.

### MICU1 and TMBIM5 exhibit reciprocal effects on mitochondrial morphology and submitochondrial localization

We next investigated whether TMBIM5 can mitigate changes observed in MICU1 knockout and *vice versa* to establish whether the two proteins work cooperatively and to clarify which protein is upstream and which is downstream. We found that overexpression of MICU1-GFP rescued the rounder shape quantified by form factor and aspect ratio (Figure 5A) of *TMBIM5* KO cells. Overexpression of TMBIM5-GFP or a channel pore mutant D294R/D325R (DM, described in (Zhang *et al*, 2022)), in contrast, had no effect on the roundish shape of mitochondria in *MICU1* KO cells (Figure 5B). A similar pattern emerged when we studied mitochondrial size. MICU1-GFP again rescued the smaller size of *TMBIM5* KO mitochondria (Figure 5C), while both TMBIM5-GFP and DM-TMBIM5-GFP surprisingly even further reduced the smaller size of MICU1 KO cells (Figure 5D). In wildtype mitochondria, only DM-TMBIM5 reduced the size of mitochondria, while TMBIM5 had no effect (Figure 5D).

**Figure 5.**
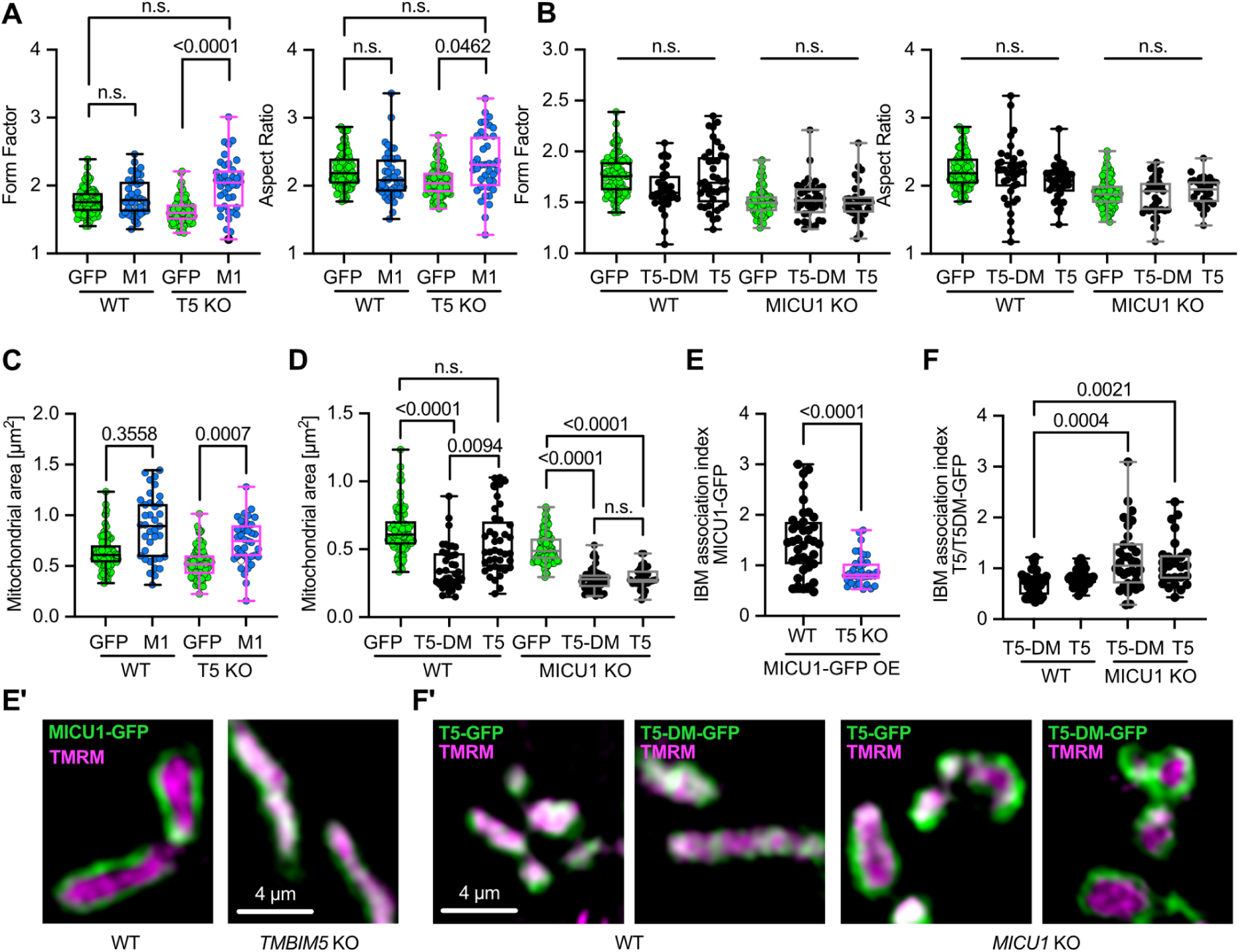
MICU1 and TMBIM5 exhibit reciprocal effects on mitochondrial morphology and submitochondrial localization. **(A)** Morphometric analysis of TMRM labeled mitochondria in WT and T5KO HEK cells expressing control vector (GFP) or MICU1-GFP (M1) imaged with structured illumination microscopy **(B)** Morphometric analysis of TMRM labeled mitochondria in WT and M1KO HEK cells expressing control vector (GFP), TMBIM5-GFP (T5), or TMBIM5DM-GFP (T5DM) imaged with structured illumination microscopy **(C)** Analysis of mitochondrial area using TMRM labelling in WT and T5KO HEK cells expressing control vector (GFP) or MICU1-GFP (M1) imaged with structured illumination microscopy. **(D)** Analysis of mitochondrial area using TMRM labelling in WT and M1KO HEK cells expressing control vector (GFP), TMBIM5-GFP (T5), or TMBIM5DM-GFP (T5DM) imaged with structured illumination microscopy **(E/E’)** Analysis of the submitochondrial localization of MICU1-GFP using the IBM association factor with TMRM as reference label in WT and T5KO HEK cells. Representative images of TMRM stained WT and T5KO HEK cells expressing MICU1-GFP (M1) imaged with structured illumination microscopy **(F/F’)** Analysis of the submitochondrial localization of TMBIM5-GFP and TMBIM5DM-GFP using the IBM association factor with TMRM as reference label in WT and M1KO HEK cells. Representative images of TMRM stained WT and M1KO HEK cells expressing TMBIM5-GFP (T5-GFP) or TMBIM5DM-GFP (T5DM-GFP) imaged with structured illumination microscopy. Data are presented as box and whisker plots with the box representing the interquartile range, spanning from the 25th to the 75th percentile. A horizontal line within the box indicates the median. Whiskers extend from the minimum to the maximum data points. Each data point represents a single cell accumulated from 4 independent experiments. Statistical significance in A, B, C, D and F was determined using the Kruskal-Wallis test followed by Dunn’s multiple comparisons test and in E using unpaired *t* test, *p* values are indicated.

When examining submitochondrial localization using the IBM association index methodology (Gottschalk *et al*, 2024), we observed patterns that contrasted with the morphological effects. Overexpression of MICU1-GFP in wildtype cells resulted in a highly positive association index, indicating predominant localization at the inner boundary membrane as described (Gottschalk *et al*, 2019). However, this spatial organization was significantly disrupted in TMBIM5 KO cells (Figure 5E), despite MICU1-GFP’s ability to rescue mitochondrial size in this background. Conversely, TMBIM5-GFP and DM-TMBIM5-GFP showed no positive correlation with the IBM in wildtype cells, but both variants shifted to the inner boundary membrane in MICU1 KO cells (Figure 5F), where they exacerbated size reduction.

These opposing patterns reveal a striking inverse correlation between mitochondrial membrane compartmentalization and morphological regulation. In TMBIM5 KO cells, MICU1 results in more prominent mitochondria despite lower association with the IBM. In MICU1 KO cells, TMBIM5 results in smaller mitochondria while showing a higher association with the IBM, irrespective of its channel function. Together, these findings suggest that TMBIM5 functions upstream of MICU1 in the regulation of mitochondrial morphology, as MICU1 can rescue phenotypes caused by TMBIM5 deficiency. The channel pore mutant DM-TMBIM5 exhibits dominant-negative effects by reducing mitochondrial size in wild-type cells, whereas wild-type TMBIM5 does not. However, in MICU1 KO cells, both wild-type and mutant TMBIM5 reduce mitochondrial size similarly, indicating that MICU1 may normally suppress a channel-independent, size-reducing function of TMBIM5 that becomes dysregulated when MICU1 is absent.

## Discussion

In this study, we characterized the interaction between TMBIM5 – a bidirectional pH-sensitive Ca²⁺ transport system in the inner mitochondrial membrane – and components of the mitochondrial calcium uniporter complex. Genetic studies in Drosophila melanogaster revealed that partial MICU1 depletion selectively rescued phenotypes in Tmbim5-deficient flies, which otherwise display disrupted mitochondrial structure and function similar to findings observed in mice. In mammalian cells, TMBIM5 and MICU1 were found in the same macromolecular complexes, and their deficiency results in comparable alterations in mitochondrial morphology and membrane potential compartmentalization. Complementation assays demonstrated that MICU1 expression restored mitochondrial morphology in TMBIM5-knockout cells, while TMBIM5 overexpression exacerbated mitochondrial defects in MICU1-deficient cells, with both proteins exhibiting opposing effects on submitochondrial distribution. Together, this suggests a functional interplay between TMBIM5 and MICU1 that maintains mitochondrial structural and functional homeostasis.

There are two potential hypotheses how partial MICU1 depletion ameliorates TMBIM5-deficiency phenotypes. Either MICU1 is directly involved in the mechanism that leads to TMBIM5-dependent mitochondrial dysfunction, or partial MICU1 depletion triggers a cellular stress response program that also protects against TMBIM5 deficiency. A direct involvement could theoretically involve MICU1’s effect on mitochondrial Ca^2+^ entry. MICU1 deficiency results in increased mitochondrial Ca^2+^ levels because loss of its inhibitory “gatekeeping” function (Mallilankaraman *et al*, 2012) allows constitutive Ca^2+^ leak into the mitochondria even at resting/low cytosolic Ca^2+^ concentrations, resulting in increased mitochondrial Ca^2+^ levels. It is unlikely that a combined MICU1 and TMBIM5 deficiency would ameliorate these potentially detrimental Ca^2+^ levels as TMBIM5 itself is also involved in mitochondrial efflux (Austin *et al*, 2022; Patron *et al*, 2022). Also, double deficiency of TMBIM5 with MCU and EMRE, which lowered mitochondrial Ca^2+^ levels, was detrimental. MICU1 has also been reported to be involved in cold-induced ferroptosis (Nakamura *et al*, 2021), a form of regulated cell death mediated by iron-dependent lipid peroxidation (Berndt *et al*, 2024). Importantly, MICU1 functions as a critical regulator in ferroptotic cell death pathways through mechanisms distinct from its canonical Ca^2+^ gatekeeping role by preventing mitochondrial hyperpolarization. A structure-function analysis revealed that domains involved in MICU1 dimerization, rather than the Ca^2+^-sensing EF-hand domains, were essential for this process, indicating a primarily structural contribution to ferroptosis regulation. In our TMBIM5-deficient model, we observed a constellation of mitochondrial abnormalities — disrupted cristae architecture, organelle swelling, and enhanced susceptibility to permeability transition — that collectively create cellular conditions conducive to ferroptotic processes. It is, therefore reasonable that partial MICU1 reduction could attenuate these pathological effects by specifically preventing the mitochondrial hyperpolarization established as a prerequisite for lipid ROS accumulation during ferroptosis, thereby providing a mechanistic explanation for the rescue phenotype observed in our double-deficient model. In addition, an involvement of TMBIM5 in hyperpolarization-triggered ferroptosis mediated by MICU1 is in line with the findings described by Patron et al. that persistent mitochondrial hyperpolarization initiates a regulatory cascade beginning with TMBIM5 degradation (Patron *et al*, 2022). This degradation subsequently releases the mitochondrial m-AAA protease AFG3L2 from inhibition, enabling its proteolytic activity. The unleashed AFG3L2 then orchestrates a comprehensive remodeling of the mitochondrial proteome, particularly targeting respiratory complex I subunits. This proteolytic breakdown of complex I components serves as a compensatory mechanism to attenuate excessive hyperpolarization by limiting electron transport chain activity and reducing reactive oxygen species production - essentially creating a negative feedback loop to restore membrane potential homeostasis under conditions of persistent hyperpolarization (Patron *et al*, 2022). Alternatively, the hypothesis that MICU1 triggers a protective stress response, which also protects against TMBIM5 deficiency, appears plausible. Such a stress response could involve activation of the DELE1 (DAP3-binding cell death enhancer 1) pathway through the altered cristae structure observed in MICU1-depleted flies. In this scenario, changes in cristae architecture would activate the protease OMA1, leading to proteolytic processing of DELE1 and translocation to the cytosol, culminating in activation of the mitochondrial integrated stress response (Fessler *et al*, 2020; Guo *et al*, 2020). Interestingly, this protection is particularly evident in skeletal muscle, which parallels the tissue-specific vulnerability observed in both TMBIM5 and MICU1 deficiency models (Lin *et al*, 2024).

The paradoxical observation that TMBIM5 and MICU1 exhibit opposing effects on submitochondrial localization and mitochondrial morphology reveals complex regulatory dynamics within the inner mitochondrial membrane. Using structured illumination microscopy methodology (Gottschalk *et al*, 2024), we observed contrasting effects on homogenization of mitochondrial membrane potential and morphological rescue effects, suggesting a dissociation between localization and functional outcomes. This inverse correlation may reflect molecular adaptation through altered interaction networks. Both TMBIM5 and MICU1 interact with CHCHD2 and MIC60, components of the MICOS complex independently of the MCUC (Meng *et al*, 2017; Tomar *et al*, 2023), which might explain their presence in shared macromolecular complexes despite distinct primary functions. When one protein is absent, the other may undergo compensatory redistribution to maintain critical submitochondrial functions. The significant shift of TMBIM5 to the inner boundary membrane in MICU1-knockout cells suggests that TMBIM5 or a complex containing TMBIM5 may assume functions typically performed by MICU1 at this location, despite this redistribution cristae architecture and preventing cytochrome c release (Oka *et al*, 2008; Seitaj *et al*, 2020), functions also attributed to MICU1 (Gottschalk *et al*, 2019). This functional overlap, combined with their differential submitochondrial distribution in knockout backgrounds, suggests that spatial organization is critical for proper mitochondrial function. The inner mitochondrial membrane contains distinct lipid microdomains, and perturbation of key proteins may alter these domains, affecting both protein distribution and membrane curvature. This relationship between localization and function appears highly context-dependent, with proteins assuming different roles depending on their submitochondrial environment and available interaction partners.

A particularly intriguing mechanistic hypothesis that should be addressed in future work involves MICU1 functioning as an intermediary signal transducer between the IMS Ca²⁺ environment and TMBIM5 activity. Given that MICU1 contains Ca²⁺-sensing EF-hand domains and our demonstration that MICU1 and TMBIM5 exist within the same macromolecular complex, MICU1 could potentially confer information about IMS Ca²⁺ concentrations directly to TMBIM5. In this model, conformational changes in MICU1 induced by Ca²⁺ binding would modulate TMBIM5’s bidirectional Ca²⁺/H⁺ exchange activity through direct protein-protein interactions. The reciprocal effects observed in our complementation studies, where MICU1 expression rescues TMBIM5-knockout phenotypes while TMBIM5 exacerbates MICU1-deficient mitochondrial abnormalities, support such a hierarchical regulatory relationship. This proposed mechanism aligns with previous observations that MICU1 has functional roles beyond MCU regulation in Drosophila, as MICU1 deficiency caused lethality that could not be rescued by simultaneous MCU knockout (Tufi *et al*, 2019). Our findings extend this concept by suggesting that these MCU-independent functions of MICU1 may include direct modulation of TMBIM5-mediated Ca^2+^ transport. This would establish a sophisticated regulatory circuit for fine-tuning mitochondrial Ca²⁺ homeostasis independent of the canonical MCU-mediated uptake pathway.

In conclusion, our findings establish a functional interplay between TMBIM5 and MICU1 in maintaining mitochondrial integrity, with implications for understanding the molecular basis of mitochondrial disorders, particularly those affecting skeletal muscle.

## Resource availability

Requests for further information and resources should be directed to and will be fulfilled by the lead contact, Axel Methner.

All unique/stable reagents generated in this study are available from the lead contact without restriction.

## Acknowledgments

This work was funded by the Deutsche Forschungsgemeinschaft (DFG) to AM (ME1922/17-1). We thank Marion Silies for providing lab space.

## Author contributions

LZ, BG, AK, FD, SB, DB, LRC performed experiments and analyzed data. LZ, BG and AM designed the experiments. VG and WFG supervised and contributed ideas. LZ, BG, VG, and WFG edited the manuscript. AM conceptualized the study, obtained funding, and wrote the manuscript.

## Materials and Methods

### Denaturing immunoblotting

To obtain protein samples, cells were directly lysed in Dodecyl-β-D-maltoside-lysis buffer (DDM-lysis buffer; 50 mM HEPES, 150 mM NaCl, 0.2% DDM, 0.5 mM EGTA, 0.3 mM CaCl_2_). Mouse tissue samples were homogenized in the same buffer at 4,000x rpm, 30 s using a glass-Teflon-potter (1-3x, until homogeneous). After 30 minutes (min) solubilisation (4°C, rotating), all samples were centrifuged (21,000x g, 10 min, 4°C) and the supernatant was used for western blotting. Protein samples were denatured in 1x Laemmli-β-mercaptoethanol-buffer, 95°C, 5 min. After gel electrophoretic separation of the proteins, they were transferred to nitrocellulose membranes by using a semi-dry blotting system (Bio-rad). For the quantification of TMBIM5, membranes were incubated in SDS-β-mercaptoethanol solution (100 mM β-mercaptoethanol, 2% SDS, 62.5 mM Tris-HCl, pH 6.7) at 55°C for 15 min on a shaker. After washing (TBST) and blocking (3% milk powder in TBST, 1 hour (h), room temperature (RT)), the membranes were incubated with the respective primary antibodies (overnight, 4°C, rotating). Antibodies used: rabbit anti-TMBIM5 (Proteintech, 1:1,000), rabbit anti-MCU (Millipore Sigma, 1:500), rabbit polyclonal anti-MICU1 (Merck, 1:500), rabbit polyclonal anti-MICU2 (Abcam, 1:500), rabbit polyclonal anti-alpha-LETM1 (Thermo Fisher, 1:500), rabbit polyclonal anti-EMRE (Santa Cruz, 1:200), rabbit anti-HA (Abcam, 1:500), mouse anti-actin (Merck chemicals, 1:1,000). Fluorescence-labelled secondary antibodies were used, and the signal was detected using a Li-Cor Odyssey imaging system and quantified with the Image Studio Lite software. The intensity was normalized to the loading control and the mean per blot.

### Blue native polyacrylamide gel electrophoresis (BN PAGE)

Protein samples were solubilised with 5 % digitonin on ice for 15 min followed by centrifugation (20,000x g, 30 min, 4°C). 0.25 % of G-250 was added to the supernatant and complexes were separated via 4-16 % Bis-Tris gels (NativePAGE ™, Thermo Fisher). The complexes were transferred to PVDF-membranes via wet blotting without methanol. Following fixation (8 % acetic acid), destaining (50 % methanol and 25 % acetic acid), and blocking (3% milk powder in TBST, 1 h, RT), the membranes were incubated with the respective primary antibodies (overnight, 4°C, rotating). The used antibodies are listed above. The staining of HRP-coupled secondary antibodies was detected with the Clarity ™ Western ECL Kit (Bio-Rad)/SuperSignal™ West Femto Maximum sensitivity Kit (Thermo Fisher Scientific) using a Bio-Rad detection system.

### Fly Stocks

*Tmbim5*/*CG2076* KO flies were obtained from Wellgenetics, Taiwan. *MCU*, *EMRE, MICU1*, and *MICU3* KO flies were provided by Alex Whitworth of the MRC Mitochondrial Biology Unit, Cambridge, UK. All flies were maintained at 25°C on standard food.

### Lifespan

Fruit flies were provided with standard molasses-based food and accommodated in a climate chamber at a temperature of 25°C with a 12 h light and 12 h darkness schedule. The flies were given fresh food every two days and dead flies were scored. Each experiment consisted of at least four groups of 25 flies in each group.

### Climbing

Climbing assays were conducted on at least four groups of 25 flies per genotype in 2 cm diameter vials, recording the percentage of flies climbing 6 cm in 10 seconds (s). Tests were standardized for time of day without CO_2_ anesthesia 24 h prior.

### Wing phenotype

Abnormal wings were scored in six-week-old flies across five groups per genotype, with each group containing 25 flies.

### ATP content

ATP levels in flies were measured following a method described previously (Liu & Lu, 2010). Briefly, two heads, thoraxes, or abdomens were lysed in 100 µl of lysis buffer from the ATP Bioluminescence Assay Kit HS II (Merck 11699709001), heated at 95°C for 2 min, and centrifuged at maximum speed at 4°C for 1 min. The assay mixed 2.5 µl of the clear supernatant, 187.5 µl of dilution buffer, and 10 µl of luciferase from the kit, and luminescence was immediately measured using a Spark Multimode Microplate Reader (Tecan). ATP concentrations were quantified using an ATP standard curve and normalized to protein concentrations measured via the BC Assay.

### Quantification of mitochondrial and nuclear DNA

Total DNA, including mtDNA, was purified using the Quick-gDNA Miniprep Kit (ZYMO RESEARCH, D3025), designed for isolating various DNA types such as genomic and mitochondrial. The mtDNA copy number was estimated by comparing the mtCOI mitochondrial gene to the nuclear gene nCOX5A, both encoding proteins involved in the electron transport chain. Detection was performed using the Faststart Universal SYBR Green Master kit (Sigma-Aldrich 4913850001) with each reaction containing 10 ng of DNA and 10 pmol of each primer. mtDNA levels were quantified using the 2^−ΔΔCt^ method, where ΔCt = Ct_mtCOI - Ct_nCOX5A and ΔΔCt = ΔCt experimental group - mean ΔCt control group.

### Functional mitochondrial preparation

Functional mitochondrial preparation followed published protocols (Tufi *et al*, 2019; Frezza *et al*, 2007). Briefly, 100 flies were homogenized in 2 ml of Mannitol-Sucrose (MS) buffer (containing 225 mM mannitol, 75 mM sucrose, 5 mM HEPES, and 0.1 mM EGTA/Tris, pH adjusted to 7.4 with 10 M KOH, all stored at 4°C) supplemented with 1% bovine serum albumin (BSA). The homogenization was conducted using a Dounce glass potter with a loose-fitting glass pestle at a speed setting of 1.5 for 15 strokes. The homogenate was centrifuged at 1,000x g for 10 min at 4°C, and the supernatant was filtered through a fine mesh (100 µm). This filtrate was then centrifuged twice at 6,000x g for 10 min each at 4°C in 5 ml of MS buffer containing BSA. The resultant pellet was resuspended in 1.5 ml of incubation buffer (10 mM Tris/MOPs, 10 μM EGTA/Tris, 5 mM Pi/Tris, 5 mM glutamate, 2.5 mM malate, and 250 mM sucrose, pH adjusted to 7.4 with HCl, all stored at 4°C) and centrifuged at 7,000x g for 10 min at 4°C. The final pellet was resuspended in 60 µl of the same incubation buffer. Protein concentration was then assessed using a BCA assay with a 1:100 dilution.

### Citrate synthase activity

Citrate synthase activity was assessed in whole protein lysates. The assay was performed in 96 well plates as follows: 2.5 µl of triton 10%, plus 2.5 µl of acetyl CoA 12.2 mM, plus 10 µl of DTNB (5,5’-dithiobis-(2-nitrobenzoic acid) 1 mM were added to each well. Then, 2 µl of protein lysate was added together with 78 µl of dH_2_O. The plate was placed in a TECAN Infinite Pro 200 reader, and the baseline acquisition was started. The absorbance was measured at OD 412 nm. After baseline acquisition, 5 µl of oxaloacetate (10 mM, pH 8.0) was added quickly to each well, and the instrument immediately started the absorbance measurements in kinetic mode at RT for 3 min in 10 s intervals.

### Transmission Electron Microscopy

Fly thoraxes were fixed using a solution containing 0.2 M cacodylate buffer (Caco-buffer), 25% glutaraldehyde (GA), and 15% paraformaldehyde (PFA). Post-fixation, tissues were washed in a 0.1 M Caco-buffer solution diluted with ddH_2_O and then incubated in 2% osmium tetroxide with 0.2 M buffer. After a second wash, samples were dehydrated in an ethanol gradient (30% to 70%) and stored overnight at 4°C. Dehydration continued the following day using 80%, 90%, 95%, and 100% ethanol, followed by washing with propylene oxide and overnight incubation in a 1:1 mixture of resin and propylene oxide. The next day, tissues were transferred to pure resin, embedded in molds, and polymerized in an oven for two days. The prepared samples were sectioned, stained for contrast, and imaged using a Tecnai 12 Transmission Electron Microscope.

### Mitochondrial PEG shrinking assay

Functional mitochondria were resuspended in assay buffer (125 mM KCl, 10 mM HEPES, 2 mM MgCl_2_, 2 mM K_2_HPO4, pH 7.2 adjusted with KOH), freshly supplemented with 100 mM succinate and 0.2 µM thapsigargin. Following a 2-min baseline measurement, polyethylene glycol-3350 (PEG, 5% final concentration) was added to induce shrinkage in pre-swollen mitochondria, requiring a large volume (100 µl) due to PEG’s high viscosity. Measurements were normalized to the final min post-PEG addition to account for baseline variations. Absorbance was recorded at 540 nm using a Spark Multimode Microplate Reader.

### Quantitative RT-PCR

Total RNA from male flies was extracted using the ZR RNA MiniPrep kit (ZYMO RESEARCH), and cDNA was synthesized from 10 ng/µl RNA using the High Capacity cDNA Reverse Transcription Kits (Life Tech). Quantitative PCR (qPCR) was performed with FastStart Universal SYBR Green Master (Rox) (Merck), employing primers sourced from Eurofins. Transcriptional levels were quantified using the 2^−ΔΔCt^ method, where ΔΔCt = ΔCt of the experimental group minus the mean ΔCt of control groups. ΔCt was calculated as Ct of the gene of interest minus Ct of the housekeeping genes RpL32/Rp49, which served as normalization controls.

### Cell culture

Cells were washed once with loading-buffer containing in mM: 2 CaCl_2_, 135 NaCl, 5 KCl, 1 MgCl_2_, 1 HEPES,2.6 NaHCO_3_, 0.44 KH_2_PO_4_, 0.34 Na_2_HPO_4_, 10 D-glucose (Carl Roth, Karlsruhe, Germany), 0.1% vitamins, 0.2% essential amino acids and 1% penicillin/streptomycin at pH 7.4. Cells were incubated in loading-buffer containing 81, 40.5, 13.5, 5.4, 2.7 or 1.35 nM TMRM (tetramethylrhodamine methyl ester, Invitrogen™) and 500 nM MitoTracker™ Green FM for 60 min. As TMRM might degrade over time in storage, TMRM concentrations were measured regularly as described elsewhere (Gottschalk *et al*, 2024).

### Structured illumination microscopy

Single and dual camera SIM imaging. The SIM setup used is composed of a 405 nm, 488 nm, 515 nm, 532 nm and a 561 nm excitation laser introduced at the back focal plane inside the SIM-box with a multimodal optical fiber. For super-resolution, a CFI SR Apochromat TIRF 100x-oil (NA 1.49) objective was mounted on a Nikon-Structured Illumination Microscopy (N-SIM®, Nikon, Austria) System with standard wide field and SIM filtersets and equipped with two Andor iXon3® EMCCD cameras mounted to a Two Camera Imaging Adapter (NikonAustria, Vienna, Austria). At the bottom port a third CCD-camera (CoolSNAP HQ2, Photometrics, Tucson,USA) is mounted for wide-field imaging. For calibration and reconstruction of SIM images the Nikon software (NIS-Elements AR 4.51.00 64-bit, Nikon, Austria) was used. Reconstruction was permanently performed with the same robust setting to avoid artefact generation and ensured reproducibility with a small loss of resolution of 10% compared to most sensitive and rigorous reconstruction settings. Microscopy setup adjustments were done as described elsewhere (Gottschalk *et al*, 2019).

### IBM Association index

The IBM association factor of TMRM was calculated as described elsewhere (Gottschalk *et al*, 2024). In short, images were subjected to background subtraction (Mosaic Suite, background subtractor, NIH) with a sliding rectangle diameter of 50 pixel. The reference channel (MTG, TMRM) was Otsu (Otsu, 1979) auto thresholded and further dilated and eroded in two independent subsets. 1 erosion and 2 dilation iterations were used. Pixel-wise subtraction of the erosion reference of the dilated reference image yields in a hollow structure, used as a mask to measure the mean intensity in the mitochondrial periphery or IBM related area in the object channel. The erosion reference served as a mask to measure the bulk or cristae mean fluorescence intensity. The ratio of IBM/CM mean intensity is a value to estimate changes of the object label distribution inside a mitochondrion, which is referred to as the IBM association index. The higher the ratio value the higher the distribution of protein label in the IBM. For image analysis, the freeware program ImageJ was used (Schindelin *et al*, 2012).

### Mitochondrial Morphology

Single 3D-SIM and time-lapsed images of TMRM were used for morphological analysis. Images were background corrected with an ImageJ Plugin (Mosaic Suite, background subtractor, NIH) and a binarization was done using a Yen auto threshold (Yen *et al*, 1995). The ImageJ particle analyzer was used to extract the mitochondrial count (c), area (a), perimeter (p), minor (x) and major (y) axes of the mitochondria. Aspect ratio (AR) was determined as:

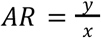

The form factor (FF) was determined as follows:

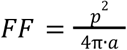

### Resources

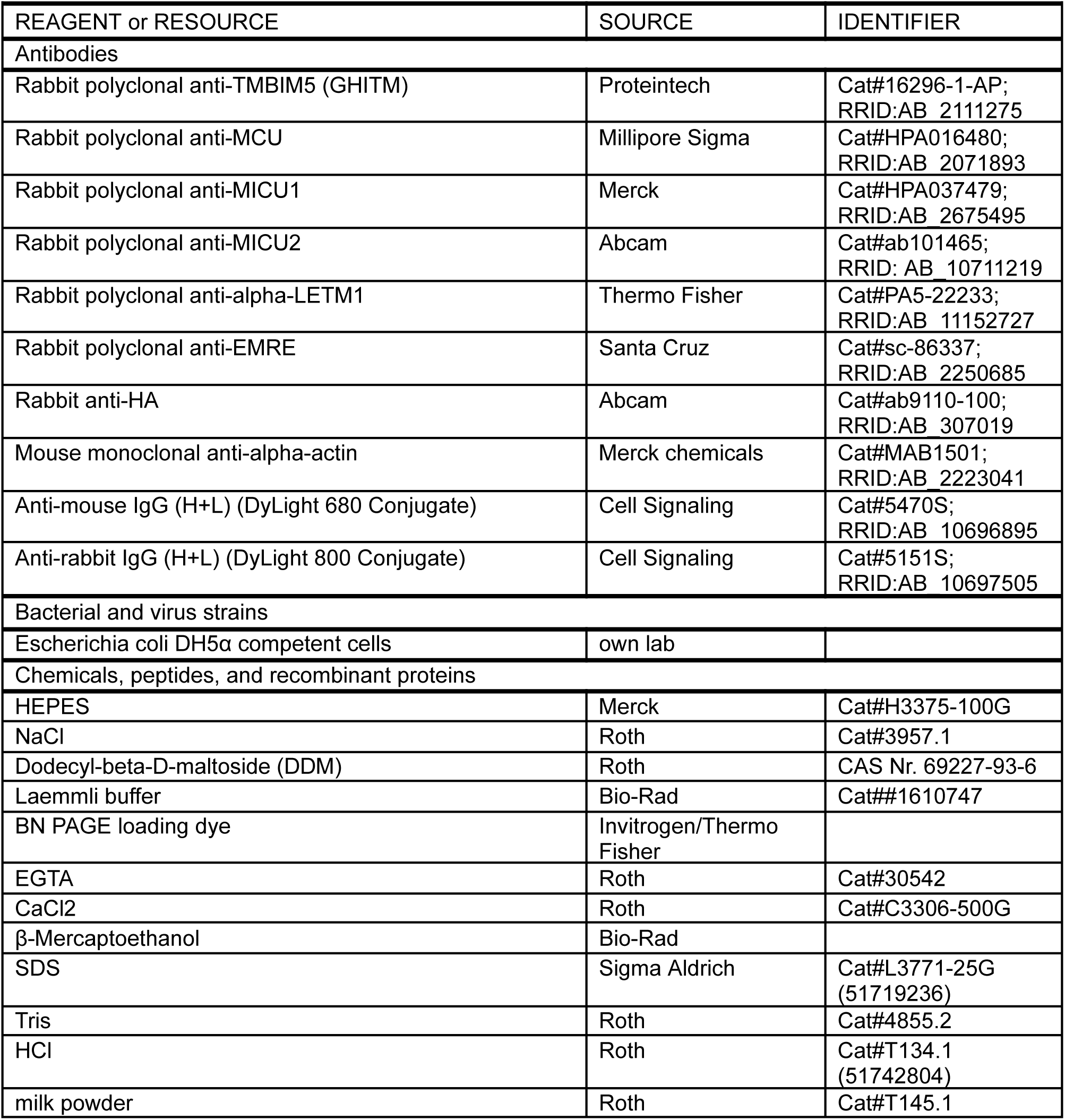
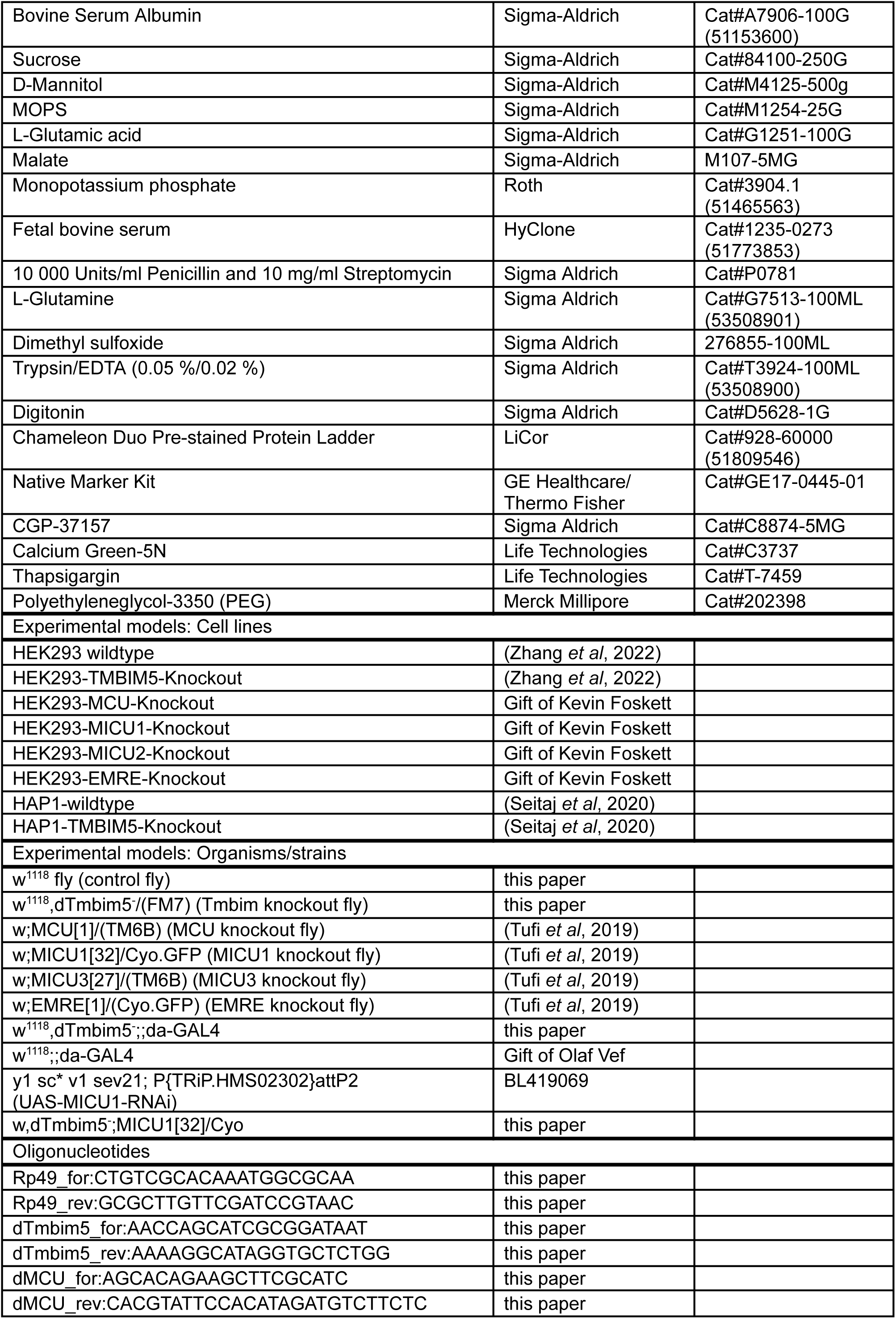
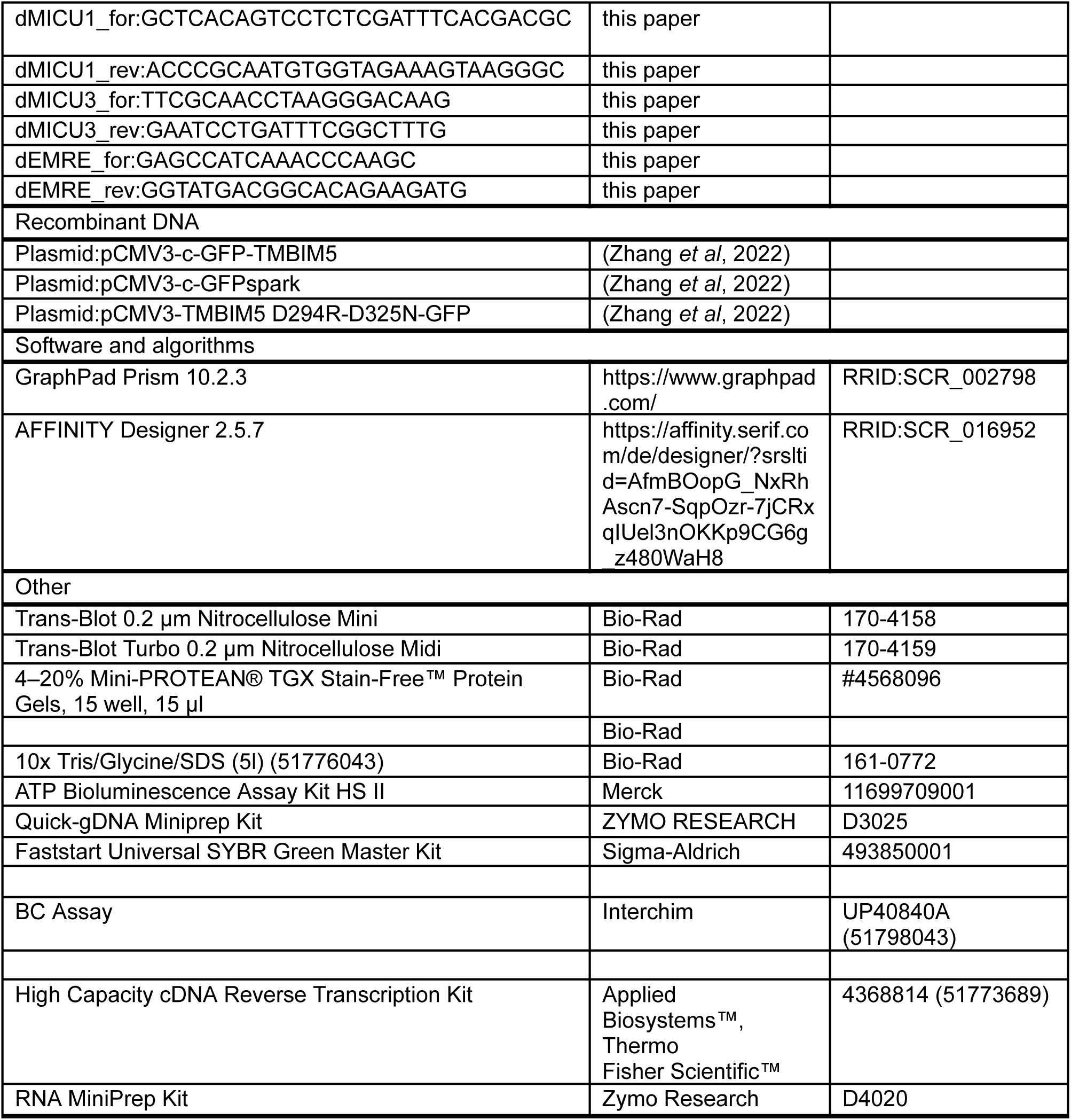

## Supplemental Figures

**Figure S1.**
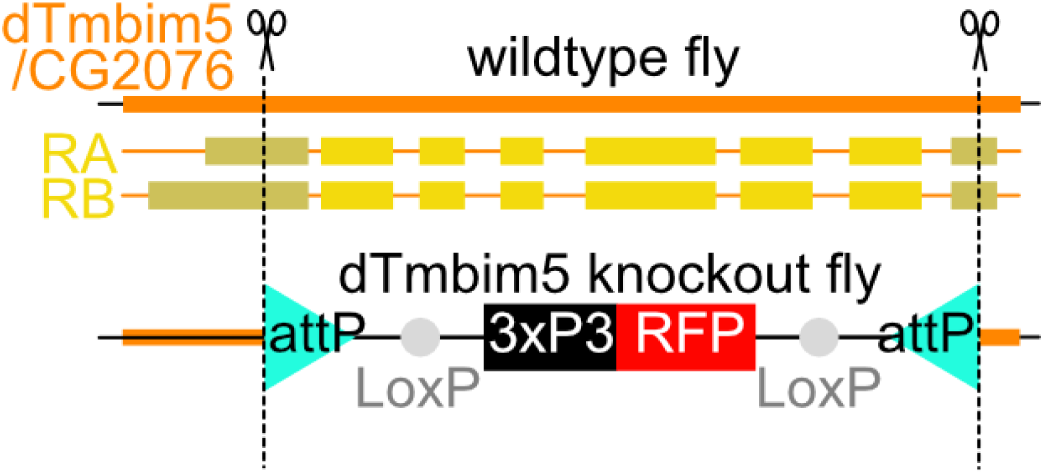
Generation of *dTmbim5* knockout flies. Knockout *dTmbim5* (*CG2076*) flies were generated using CRISPR/Cas9-mediated homology-directed repair deleting a 1,460-bp fragment (−36 nt to +1,424 nt from the ATG of *CG2076*) and replacing with a 3xP3-RFP cassette flanked by attP sites. RA and RB indicate two different transcript variants.

**Figure S2.**
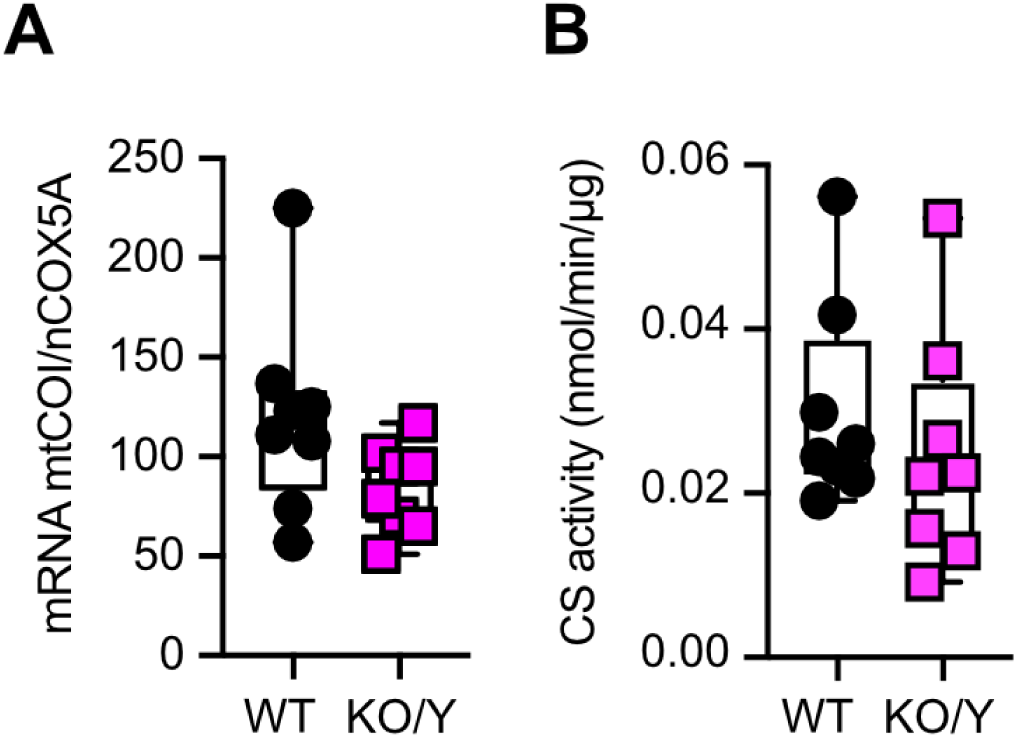
Assessment of mitochondrial content in *Tmbim5* knockout flies reveals no significant alterations in mitochondrial mass. (A) Mitochondrial DNA copy number was quantified by comparing the relative abundance of mitochondrially encoded *mtCOI* (Drosophila ortholog of human cytochrome c oxidase subunit I) to nuclear-encoded *nCOX5A* (Drosophila ortholog of human cytochrome c oxidase subunit 5A) using quantitative PCR analysis of total DNA extracts. (B) Citrate synthase (CS) activity was measured in mitochondrial preparations. Citrate synthase is a nuclear-encoded enzyme localized in the mitochondrial matrix whose activity remains stable under various metabolic conditions, making it an ideal reference enzyme for normalizing mitochondrial content across experimental conditions. Data are represented as box and whisker plots showing the interquartile range (25th to 75th percentile) with the median indicated by a horizontal line. Whiskers extend from minimum to maximum values. WT represents wild-type (*w^1118^*) flies, while KO/Y indicates hemizygous *Tmbim5* knockout males. Statistical analysis was performed using unpaired t-tests which revealed no significant changes.

**Figure S3.**
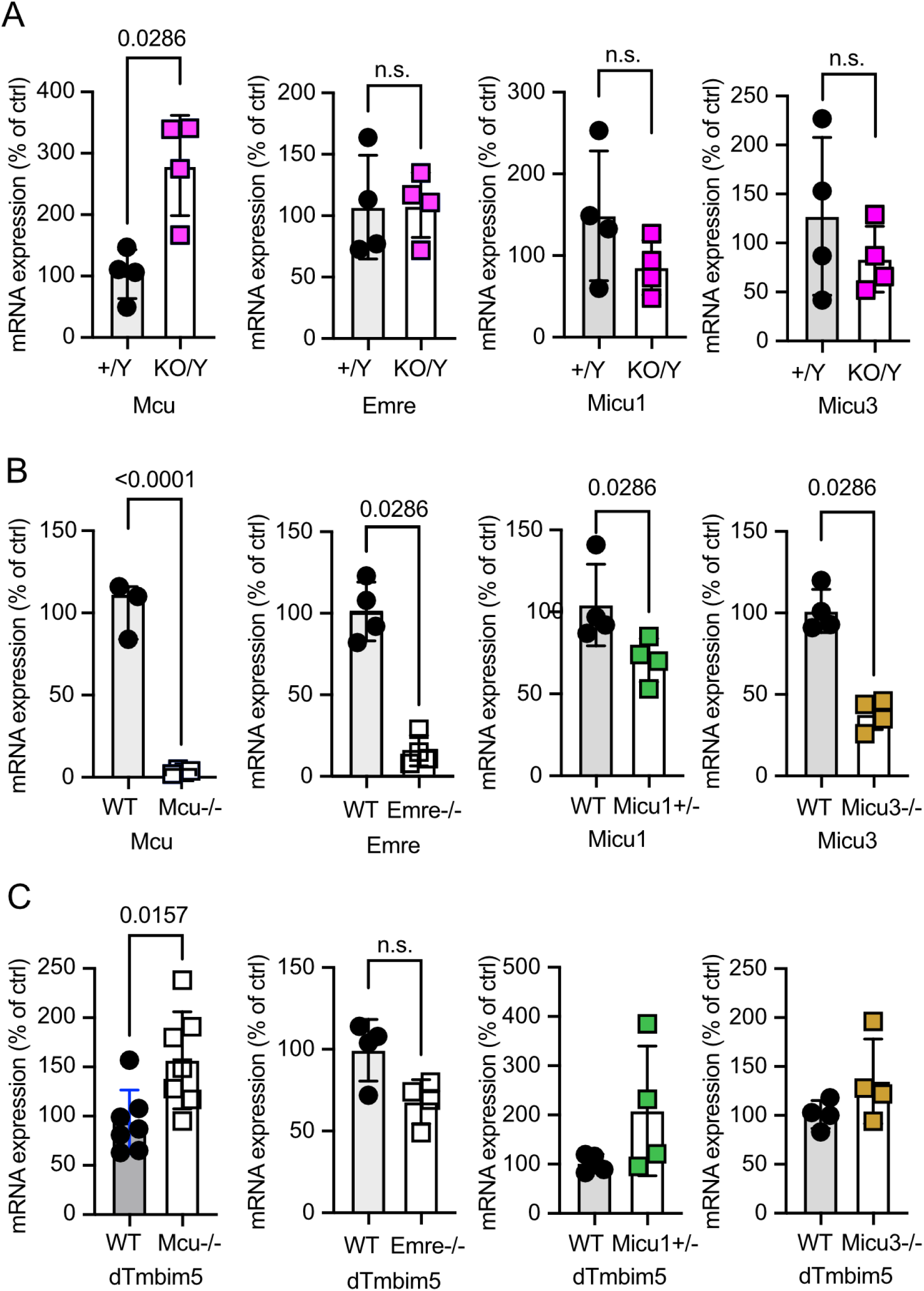
Reciprocal transcriptional regulation between *Tmbim5* and *Mcu* in Drosophila melanogaster. **(A)** Quantitative PCR analysis of mitochondrial calcium uniporter complex components in *Tmbim5* knockout (KO/Y) male flies compared to wildtype (+/Y) controls. Data demonstrate significant upregulation of *Mcu* mRNA levels while *Emre*, *Micu1*, and *Micu3* transcript abundance remains unchanged. Expression values were normalized to *Rp49* as an endogenous reference gene. (B) Validation of genetic models through transcript quantification in respective knockout lines: *Mcu* knockout (*Mcu-/-*), *Emre* knockout (*Emre-/-*), *Micu1* heterozygous knockout (*Micu1+/-*), and *Micu3* knockout (*Micu3-/-*). Quantitative PCR confirms significant reduction of target transcripts in each genetic background compared to wildtype controls. **(C)** Quantitative PCR analysis of *Tmbim5* expression across mitochondrial calcium uniporter complex mutant backgrounds. Data reveal significant upregulation of *Tmbim5* transcript levels specifically in *Mcu-/-* flies, with no significant changes observed in *Emre-/-*, *Micu1+/-*, or *Micu3-/-* genetic backgrounds. Data across all panels are presented as bar graphs showing mean ± standard deviation. Each data point represents the mean value derived from pooled RNA of 5 flies. Statistical significance was determined using the Mann-Whitney test with *p* values indicated.

